# Achieving Micrometer-Scale 4D X-ray tomography of Living Leaf Tissue in the Laboratory

**DOI:** 10.1101/2025.11.21.689754

**Authors:** F. Siracusa, E.V. Kristensen, A. Szameitat, D. J. Tobler, I. Ertem, M. Frank, C. Wu, R. Abdurrahmanoglu, A. Pinna, F. Minutello, S. Husted, J.-C. Grivel, R. Mokso

## Abstract

A methodology for achieving micrometer-scale 4D X-ray lab microscopy of living leaf tissue was developed to overcome challenges associated with delicate tissues, radiation damage, and motion artifacts during *in vivo* imaging. The study focused on optimizing laboratory based X-ray micro-computed tomography (microCT) parameters to balance high-resolution imaging with minimized physiological stress and radiation dose quantification. Assessing the dose-safe imaging window required comparing vertical and horizontal leaf mounting setups. Results demonstrated that the horizontal setup provided greater stability, preventing tissue degradation and maintaining sample viability during continuous acquisitions lasting up to 22 hours (∼15600 Gy). MicroCT capacities were clearly able to resolve microstructures at the cellular level, achieving a pixel size down to 1 µm. Furthermore, this optimized methodology confirmed the ability to track the spatiotemporal dynamics of applied compounds such as iohexol and aggregated nanoparticles within the leaf tissue. This work establishes that accessible laboratory based microCT enables the *in vivo* 4D monitoring of anatomical and physiological changes in living plants.

## 1. Introduction

Plant growth and development depend on fundamental structures, such as vascular tissue and stomata, which control key processes including water transport, gas exchange and nutrient distribution (Brodribb et al. 2007; McElrone et al. 2013; Sack & Holbrook 2006). Studying their architecture and responses to, for example, environmental cues can provide deeper insights into how these structures regulate essential plant functions. However, direct observation under natural, undisturbed conditions is challenging because the relevant plant tissues are delicate and can be easily damaged, or, in the case of the plant vasculature, this is not easily accessible. State-of-the-art techniques to probe plant tissue down to the micrometer scale include electron and confocal microscopy techniques (Truernit & Palauqui 2009; Moreno et al. 2006). Amongst these, confocal laser scanning microscopy (CSLM) is probably the most widely used method for non-destructive imaging of living cells in real time (Moreno et al. 2006, Pasternak et al. 2015), however, its capabilities are limited due to the depth penetration of the laser, often requiring organelle labelling with fluorescent dyes (Pinna & Husted, 2025). Electron microscopy approaches can reach the highest resolutions down to nanometer scale but require elaborate sample preparation, including fixation, staining, and/or sectioning (Deal et al. 2015; Yuan et al. 2020) which impairs the study of living organisms.

MicroCT offers a range of interesting solutions to these limitations, by providing high penetration depth and eliminates the need for physical sample sectioning and organelle labelling. Indeed, over the past two decades, X-ray microCT has gained growing interest in plant science for its ability to capture 3D internal structures without damaging plant integrity (Dhondt et al., 2010; Chen et al., 2021; Piovesan et al., 2021; Frank et al. 2025; Kristensen et al. 2025, Avellan et al., 2019; Santana et al., 2020). This has enabled research exploring tissue pore space (Herremans et al., 2015; Chen et al., 2021; Gao et al., 2023), water and solute movement in roots and stems (Cochard et al., 2015; Hou et al., 2022; Camboué et al., 2024, Gargiulo L. et al. 2024, Metzner R. et al. 2022), and the measurement of crystalline formations (Earles et al., 2018). However, most of these studies focused on non-living plant samples (e.g., wood) or rigid living structures such as roots, vines, and conifer needles (Chen et al., 2021; Hou et al., 2022; Gao et al., 2023), as well as detached parts like buds, fruits, seeds, and leaves (Mathers et al., 2018; Duncan et al., 2022), (SI Table S1).

Among the plant organs, leaves are particularly challenging to study *in vivo* with X-ray microCT due to their thin, delicate tissues and dynamic physiological processes, which make them highly susceptible to radiation damage and prone to motion artifacts. Extended X-ray exposure can lead to shrinkage or cell collapse, while tissue movement may distort 3D image reconstruction. Increasing the field of view (FOV) - the portion of the leaf being imaged - often involves a trade-off with resolution, yet it can improve the success of *in vivo* imaging, as demonstrated in studies such as Pfeifer et al. (2018). However, in these cases, the relatively low resolution (pixel size, (PX) < 50 µm), has limited the ability to resolve fine leaf microstructures.

In our previous work, we demonstrated a new synchrotron based microCT imaging technique, which enabled studying foliar water films on live and undamaged barley and potato plants in a controlled atmosphere over time (Frank et al. 2025). By exposing plants to low CO_2_ and high humidity to keep stomata open, optimal conditions were created to activate cuticular and stomatal water uptake pathways. In this study, we use a similar sample setup and explore the limits of laboratory (lab-) based microCT for *in vivo* imaging of plant microstructure. Notably lab-based X-ray microCT is more easily accessible, flexible and user friendly, compared to synchrotron based microCT, but it is limited by lower beam coherence (resulting in reduced phase contrast), by its polychromatic energy spectrum and finally by resolution and X-ray flux. This in turn means longer acquisition times, making it crucial to maintain plant health during several hours of imaging within an X-ray shielding dark scanner. The specific objectives were thus to assess the X-ray dose-safe imaging window and the resolution limits for in vivo leaf imaging using lab-based microCT, including comparison between horizontal versus vertical leaf studies. Additionally, the study aimed to assess if surface-applied compounds can be imaged and tracked in living leaves using lab-based microCT. The physiological activity of *in vivo* microCT imaged leaves was monitored by performing photosynthesis measurements, and leaf structural changes, including shrinkage, were evaluated from the acquired microCT data. The possibility to trace the mobility of nanoparticles (NPs) and/or liquids when applied to plant leaves was tested using gold, manganese dioxide and hydroxyapatite NPs as well as iohexol, a contrast agent used in our previous synchrotron based CT study (Frank et al. 2025). It is important to note that individual NPs are below the microCT resolution, but they can be detected once aggregated into micrometer-scale clusters, as also shown in a recent synchrotron based microCT study on plant leaves (Kristensen et al. 2025). Overall, results highlight that microCT capacities in the laboratory are clearly able to resolve leaf structures such as stomata, substomatal cavities and vascular features at the cellular level, with a pixel size (PX) down to 1µm, in living plants. Notably, here we demonstrated that by using the right experimental sample holder, plant physiological processes can be maintained during several hours of imaging, thus opening the window for in-situ 3-D monitoring of physiological and anatomical changes in living plants with lab-based microCT.

## 2. Materials and Methods

### 2.1 Plant growth and monitoring

Spring barley (*Hordeum vulgare* L. c.v. KWS Irina) was used for imaging. Seeds were germinated for seven days in vermiculite. Uniform seedlings were selected and transferred to 5 L aerated hydroponic containers supplied with nutrient solution. Plants were grown in a greenhouse under controlled conditions for seven days prior to measurements (specific growth conditions detailed in SI Text S1). For each experimental session, plants were moved from the greenhouse to the scanner and maintained for at least one hour under a full-spectrum growth lamp (Cultilite LED Antares 180W COB) to allow acclimation and promote stomatal opening. Physiological activity of selected scanned leaves area was assessed with a Ciras-3 Portable Photosynthesis System (PP Systems, USA) by measuring transpiration rate (E) and stomatal conductance (gₛ) with five replicate measurements per sample.

### 2.2 Imaging Setups

Two sample setups were designed for microCT imaging, both enabling 360° tomographic imaging with roots placed in a nutrient solution (Figure 1 A, B). In the first setup, leaves were mounted vertically (Figure 1A) to achieve high spatial resolution and precise alignment of the region of interest within a small field of view (FOV). In the second setup, leaves were positioned horizontally on Parafilm supports (∼a few millimeters in diameter), preventing them from lying completely flat and reducing the amount of tissue intersecting the beam path. This configuration is particularly suited for imaging leaves before and after solution treatment (e.g., foliar applied fertilizer or pesticide). In some cases, the setup was adapted by sealing the sample area within a chamber to enable controlled humidity and gas conditions for stomatal regulation. This configuration was used to investigate plants with forced stomatal opening (Frank et al. 2025).

**Figure 1:**
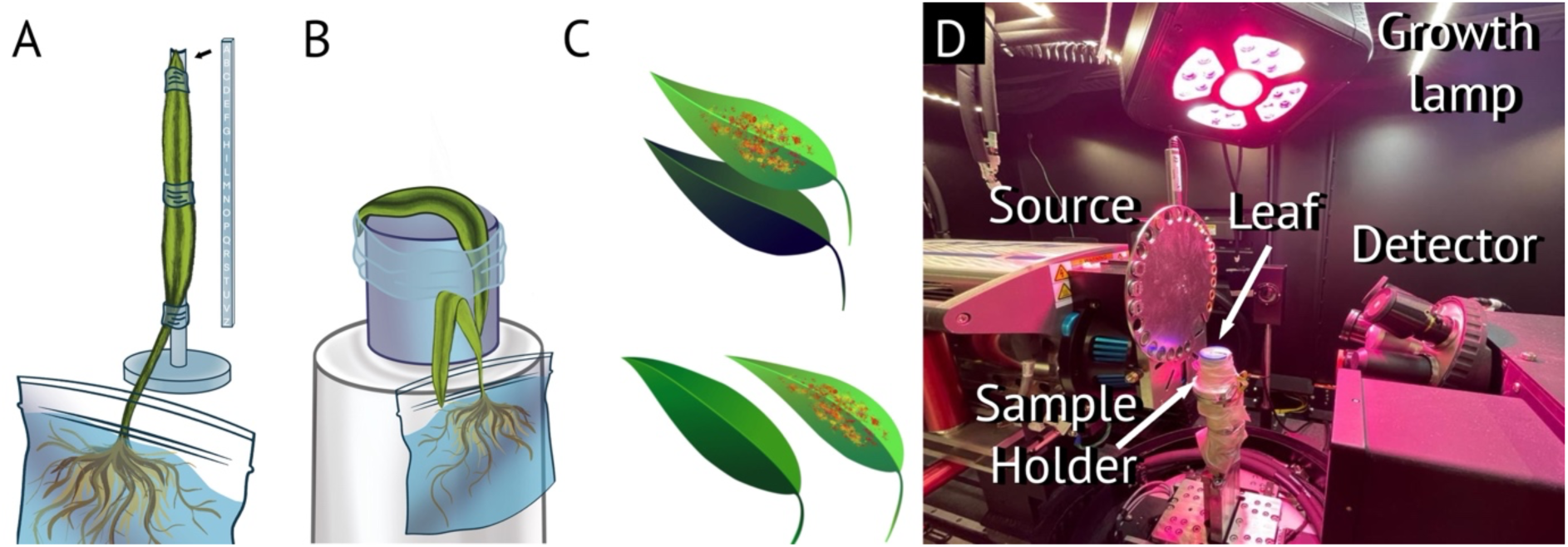
Sketches of vertical (A) and horizontal (B) leaf setups for microCT imaging. The vertical holder uses a 3D printed alphanumeric reference to track leaf spots and align scan volumes. C) Arrangement of multiple leaves, either overlapping (used only in horizontals setup) or side-by-side (both used in vertical and horizontal setup). (D) Horizontal setup with a living leaf used in the Versa microCT.

In addition to these setups, a system was developed to scan multiple leaves simultaneously, enabling direct comparison of control and treated samples. Two configurations were used: i) side-by-side for larger FOV, and ii) stacked for smaller FOV (< 3×3 mm^2^), (Figure 1C), where thin Parafilm layers separate overlapping leaves. In the stacked setup, the lower leaf receives less light, making it suitable mainly for short experiments, volume stitching, or dark-adapted plants. In contrast, the side-by-side setup exposes both leaves to identical environmental conditions. Temperature and humidity were continuously monitored near the samples using a combined hygrometer/thermometer in all the experiments. The experimental sessions consisted either in single scans or time-series where the leaves remained fixed on the sample holders for up to 22 hours.

### 2.3 Leaf treatment with NPs suspension and contrast agents

We imaged plants untreated and treated with NP suspensions or iohexol. NP suspensions were applied to leaves via two methods: i) direct infiltration, where 0.5 ml suspension were gently pushed into the leaf with a syringe to assess internal distribution, and ii) drop application, where five 5µL droplets per leaf tested surface responses (e.g., stomatal opening), penetration, and internal visibility. The following three types of NPs were tested: polyacrylic acid-coated manganese dioxide NPs (PAA-MnO_2_ NPs; Pinna et al. 2025;), hydroxyapatite NPs (nHAp; Minutello et al. 2025) and gold (Au) NPs (Burve et al. 2023), with synthesis and characterization details provided in the SI (Text S2, Figure S1). Iohexol contrast agent (150 mM, VWR, US) was applied as five drops of 10-15µl. Both iohexol and NPs suspension (except Au NPs) were supplemented with Silwet Gold (0.0025 - 0.05 %, HD 2412, DK) to enhance uniform diffusion across the leaf surface (Frank et al. 2025).

### 2.4 X-ray microCT measurements

X-ray imaging experiments were conducted with the Zeiss Versa 520 (Carl Zeiss X-ray Microscopy, Inc.) and Exciscope Polaris (Exciscope) microtomographic scanners maintaining fixed tube voltage (60 kV for Versa, 70 kV for Polaris), current (83 µA for Versa, 2642 µA for Polaris), source-to-sample distance (18 mm for Versa, 155 mm for Polaris), and number of projections (1001 for Versa and 1601 for Polaris). The Versa allows high spatial resolution with PX down to 1 µm and FOV between 1 x 1 to 8.6 x 8.6 mm^2^, whereas the Polaris is optimized for fast acquisition and larger FOV (10.63 x 7.79 mm^2^), albeit with coarser PX of 2. 5 µm and above. Scan time (ST) was adjusted according to target PX to balance image quality and acquisition time, and it varied between 25 min and 5 hours per scan (SI Table S2). Phase contrast on the Versa system was achieved by increasing the source-sample distance to ∼18 mm, which enhanced spatial coherence. We used the built-in LE1 filter to remove the low energy end of the X-ray spectrum, which is a viable approach to reduce X-ray radiation dose on the sample. The Polaris system is inherently optimized for phase contrast achieved by placing the sample at a large distance from the source (typical 150 mm) and the detector 600 mm further downstream. This geometry results in an X-ray magnification of 5.5 and a PX of 2.55 µm. Angular projections were used as input for conventional cone-beam tomographic reconstruction to generate volumetric datasets. Data obtained with the Polaris were further processed with ImageJ (Fiji) for rapid inspection, and with Dragonfly (2024.1, Comet Technologies) and Avizo (Thermo Fisher Scientific) for segmentation and volume rendering. For data obtained with the Versa, non-local-means filtering was applied in Avizo to enhance image contrast and facilitate segmentation of plant structures.

### 2.5 Dose quantification

The absorbed dose delivered to the leaf sample was calculated using the X-ray spectra of the Excillum MetalJet and Versa sources under the specified beam conditions (SI Text S3) (Excillum, 2025; Carl Zeiss X-ray Microscopy, Inc., 2025). The energy-dependent dose (Gy) was calculated (SI Figure S2), while dose gradients across the leaf surface were neglected, assuming the dose in lower-flux regions is minor relative to our estimates. The sample area was not included in the calculation, under the assumption of thin samples and uniform illumination within the source solid angle. Based on this approach, the total accumulated dose for the shortest scan on each instrument was estimated at ∼600 Gy for Polaris (25 min, PX 2.55 µm) and ∼285 Gy for Versa (45 min, PX 2.9 µm). The spectral dose profile highlights energy-dependent absorption features, with peaks corresponding to the most intense spectral components (SI Figure S2).

## 3. Results and Discussion

### 3.1 Plant response to radiation dose

Achieving high resolution with lab-based microCT requires prolonged imaging and high X-ray fluxes, making radiation damage a central concern. To evaluate the impact, data from 15 time-series microCT measurements of barley leaves (incl. between 2 and 21 scans each) were analysed, encompassing vertical and horizontal setups under various acquisition modes. The radiation dose in experiments performed here was mainly influenced by variation in the number of scans and exposure time (SI Text S3, Figure S2) as tube voltage and current, number of projections, and source-to-sample distance were kept constant for Versa and Polaris measurements, respectively (Section 2.4). Radiation-induced stress in plants was evaluated by monitoring changes in leaf thickness. These changes were quantified using the thickness ratio (*Th*), defined as the thickness given by the first scan relative to subsequent scans, and plotted against radiation exposure.

Clear differences in stress response were observed between leaves imaged in the vertical and horizontal configurations. In the vertical orientation, shrinkage and motion artifacts appeared after ∼ 5 hours of continuous scanning, corresponding to an X-ray dose ∼3500 Gy (Figure 2A), progressing to deformation and eventual collapse by 14 hours (i.e., ∼10000 Gy). In contrast, leaves imaged in the horizontal setup showed no signs of shrinkage or degradation (Figure 2A), even during the longest measurement that permitted up to 22 consecutive 1-hour acquisitions (∼15600 Gy) on the same leaf without loss of quality. To get insights into plant vitality following X-ray exposure, stomatal behavior and transpiration rates were measured in two irradiated leaves (in vertical position), one exposed to a 2-hour scan (∼1400 Gy) and one exposed to two consecutive 2-hour scans (∼2800 Gy), and then compared with an unexposed leaf kept outside the scanner. The stomatal conductance (gₛ) and transpiration rates (E) for both the 2-hour irradiated and control leaf were identical (23.8 mmol m⁻² s⁻¹, and 0.62 mmol m⁻² s⁻¹), indicating no change in vitality during the 2-hour exposure. However, for the 2 x 2-hour irradiated leaf, the gₛ and E values decreased to 12 mmol m^−2^ s^−1^ and 0.33 mmol m^−2^ s^−1^, respectively, indicating X-ray-induced damage after 4 hours of X-ray exposure. Both the irradiated plants and the control plants were then returned to the greenhouse and plant vitality was examined after 24 h. Gas exchange values on the 4-hour irradiated leaf increased once back in the greenhouse (gₛ = 33.8 mmol m⁻² s⁻¹, E = 0.8 mmol m⁻² s⁻¹), likely due to higher humidity and more controlled conditions in the greenhouse, and exhibited similar values to the 2-hour irradiated leaf (gₛ = 39 mmol m⁻² s⁻¹, E = 0.89 mmol m⁻² s⁻¹) and the control leaf (gₛ = 31.2 mmol m⁻² s⁻¹, E = 0.74 mmol m⁻² s⁻¹). However, after 48 h, the irradiated region (on both irradiated leaves) developed visible yellowing, consistent with chlorosis due to chlorophyll degradation, which became more pronounced after 4 days (Figure 2C, right).

**Figure 2:**
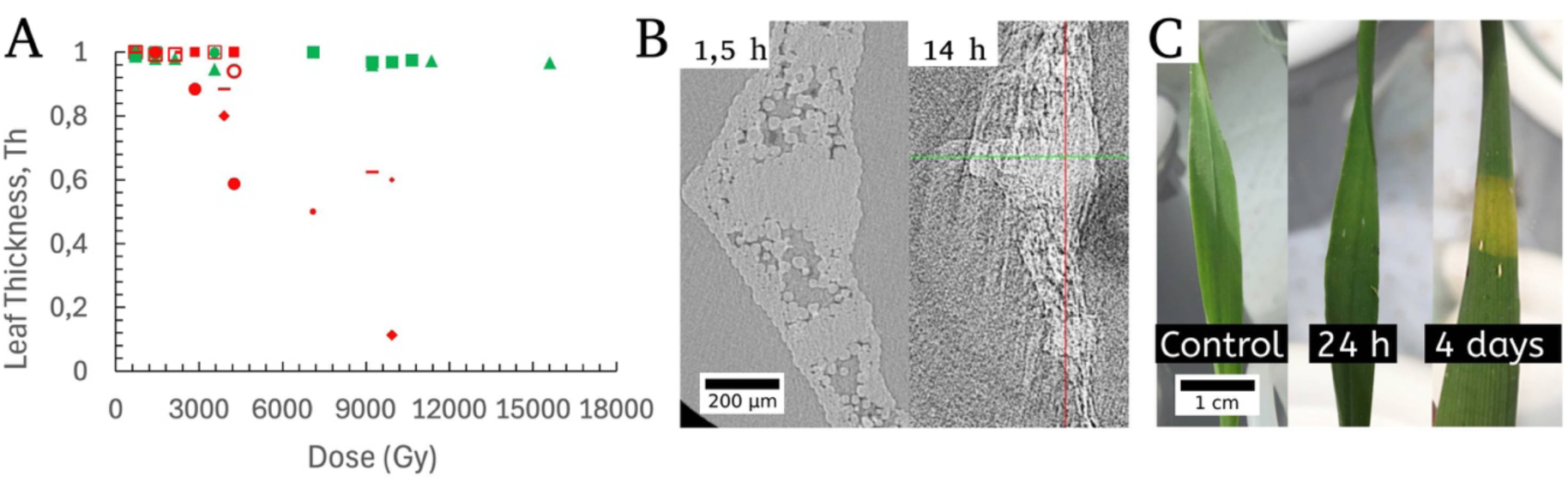
(A) Changes in leaf thickness (Th) as a function of X-ray radiation dose for experiments performed in vertical (red symbols) and horizontal (green symbols) setup. (B) Example microCT slices showing stable and collapsed leaf following 1.5 (∼1000 Gy) and 14 h (∼10000 Gy) of X-ray imaging. (C) Pictures showing stochastic radiation damage appearing 4 days following a 2 h microCT scan (∼1418 Gy). Notably, no damage is seen 24 h after the scan; control leaf with no X-ray exposure shown on the left. Radiation dose as a function of exposure time is shown in SI Figure S2. Scan details are described in SI Table S3

In summary, both setups are suitable for imaging leaves for up to 2 h with PX down to 1 um (∼1400 Gy) without inducing tissue damages, motion artefacts and structural changes, while maintaining plant vitality. The vertical orientation enables high-resolution stitching of leaf regions, but care must be taken to avoid excessive overlaps between stiches. In contrast, the horizontal setup provides greater stability for extended imaging. Its gentler immobilization and broader X-ray flux likely contribute to the reduced tissue damage observed. Additionally, the horizontal configuration allows formulations (e.g., fertilizer, pesticide) to be applied to the leaf surface between scans without disturbing the setup (discussed in section 3.3).

### 3.2 Visualisation of physiological processes at the cellular level

In our previous work, open and closed stomata were successfully imaged using synchrotron-based microCT (Frank et al., 2025). Building on this, lab microCT was employed to directly compare stomatal dynamics in treated and untreated living leaves under identical imaging conditions. To achieve this, two leaves were mounted side-by-side within the same FOV using the Polaris instrument, enabling paired observations during sequential scans (Figure 3A). An initial scan of both leaves (both untreated) showed stromata were consistently open on both the adaxial and abaxial surfaces, confirming favorable conditions for liquid uptake. Five drops of nHAp suspension were then applied to the adaxial side of one leaf, after which both leaves were scanned once more. Following application, some morphological changes could be observed: the previously open stomata on the treated adaxial surface were no longer visible, and the surface appeared continuous. In contrast, the abaxial surface of the same leaf retained open stomata (Figure 3A).

**Figure 3.**
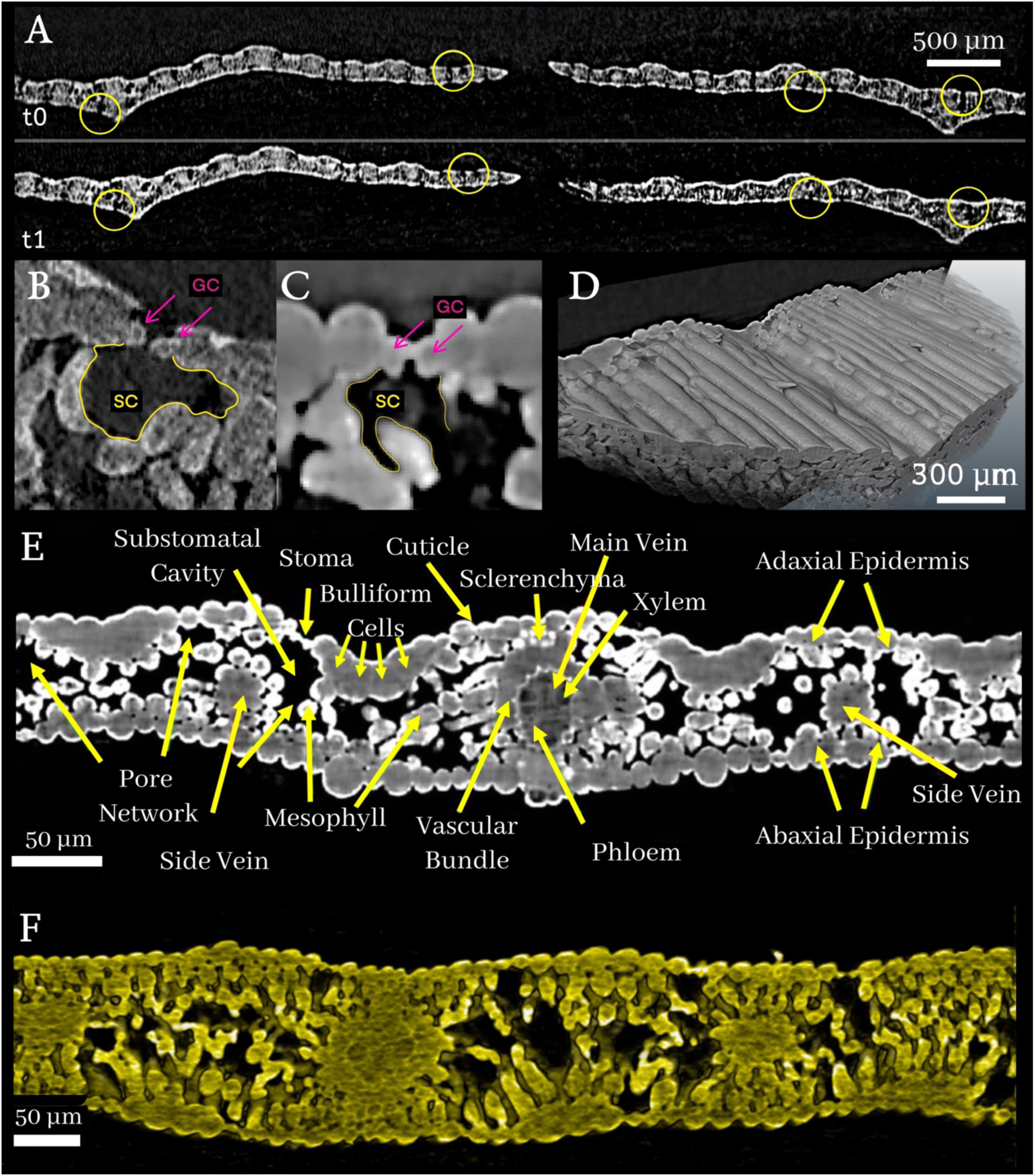
(A) Example microCT cross-sections of two barley leaves imaged side-by-side in the same FOV, before (t0) and after (t1) NP application. The leaf on the left is the control, and the leaf on the right was treated with NP. Yellow circles mark stomata locations. Close-up of microCT cross-sections showing open stoma (B) and closed stoma (C) with pink arrows pointing at guard cells (*GC*) and yellow lines delineating substomatal cavities (*SC*). (D) MicroCT volume rendering of barley leaf with visible stomata on adaxial surface. (E) MicroCT cross-sections of a grass (SI Video S1) and (F) volume rendering of imaged barley leaf (SI Video S2). Scan details are described in SI Table S3.

Comparison with the neighboring, untreated leaf after the second scan, indicated a higher number of open stomata on both adaxial and abaxial surfaces. Together, these observations could suggest that stomata on the treated adaxial surface were occluded by the applied nHAp suspension, whereas the untreated surfaces remained structurally unaffected. The key advantage of this approach is the ability to perform sequential scans of the same leaf before and after treatment without introducing motion artifacts, thereby providing greater confidence in data interpretation compared with conventional methods that rely on separate control and treated samples. Moreover, simultaneous imaging of treated and untreated leaves within the same FOV allows discrimination between structural changes induced by X-ray exposure and those specifically attributable to the applied treatment.

While an increased FOV inherently reduces spatial resolution and may limit the ability to monitor fine physiological processes, this drawback can be addressed through a two-step imaging strategy. First, larger FOV scans, performed with appropriate controls, can be used to identify regions of interest and verify sample integrity. Subsequently, selected areas can be imaged at higher resolution using smaller FOV. Applying this strategy, detailed visualization of stomatal and sub-stomatal features was achieved with the Versa scanner at 1 µm PX (Figure 3B–D). Virtual slices from the reconstructed volumes resolved guard cells and sub-stomatal cavities in both open and closed states (Figure 3B, C), while three-dimensional datasets captured the stomatal complex in situ, including cuticle, trichomes, and epidermal layers across the full leaf width. Notably, these structural details were resolved even at a very short ST of 46 min (Figure 3D). To give another example of the level of structural detail achievable, Figures 3E and 3F show denoised and phase-retrieved tomographic slices respectively of grass and barley acquired with the Versa scanner at 1 and 1.4 µm PX, and ST of 1.5 h and 5 h, respectively. A single grass *(Elytrigia)* leaf was included here as another example, illustrating the many features it shares with barely leaves. Conventional reconstruction enabled visualization of the interface between leaf tissue and air, as well as differentiation of vascular structures and distinct cell types. Although high-quality datasets could be obtained with short exposure times, the best results were achieved with scan durations between 1.5 h and 5 h, enabling resolution of vascular elements, such as phloem and xylem, which are otherwise difficult to identify.

### 3.3 Tracking applied formulations in plant leaves

Understanding how foliar-applied formulations, such as nutrients or pesticides, are taken up and distributed, remains a central question in plant leaf physiology. Here, we tested the potential of lab-based microCT to track these processes by applying NPs to barley leaves through infiltration and drop application using different configurations. The initial experiments employed Au NPs and iohexol as high contrast materials. Although neither of them represent a nutrient or growth-essential compound, they were chosen to trace infiltration pathways and assess technique feasibility.

The first experiments were made using Au NPs using infiltration to test the visibility of NPs within the leaf structure (Figure 4 A, B). Distinct clusters of Au particles were observed in treated leaves, despite the relatively low imaging resolution, whereas non-treated leaves showed no clusters (Figure 4 A). Further validation was achieved with laser ablation ICP-MS of treated leaves (fixed, sectioned and analyzed following CT; SI Figure S3), confirming the presence of gold clusters inside the tissue.

**Figure 4:**
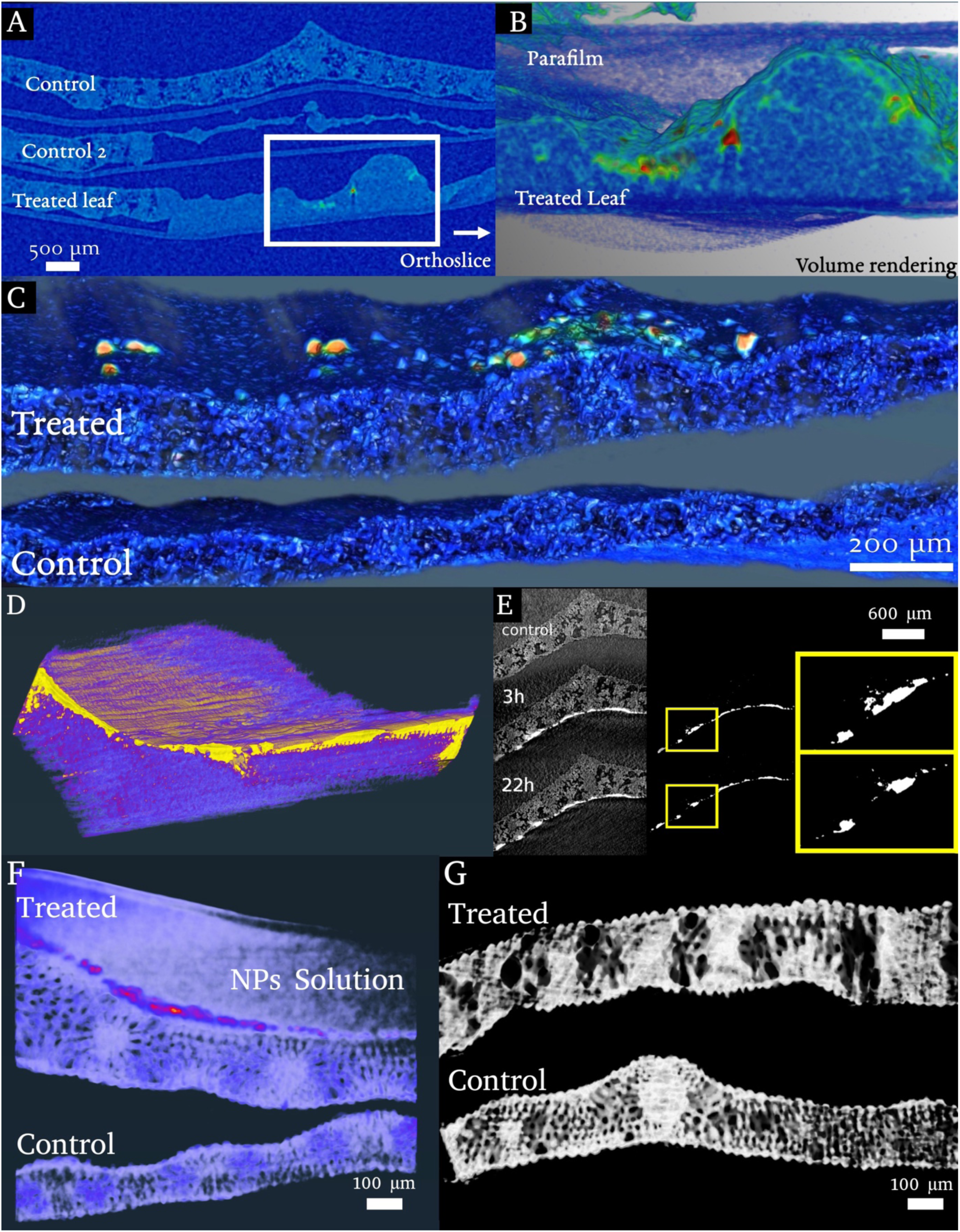
Lab-based microCT of barley leaves treated with Au NPs, iohexol and nPAA-MnO_2_ NPs. A) Orthoslice of three barley leaves: top and middle leaf are controls (middle leaf showing signs of damage due to mishandling), and bottom leaf was treated with AU NPs. B) Volume rendering of zoomed-in region indicated by the white square in A (SI Video S3). The red/green regions indicate clusters of higher density material, attributed to the accumulation of Au NPs). C) Volume rendering of two leaves: top leaf treated with Au NPs, bottom leaf is control. (D) Volume rendering of a leaf treated with iohexol (SI Video S4), scanned together with a control leaf (not in figure). E) Slice from the same leaf visible in three time-steps and visualization of the thresholded contrast media with close-ups. F) Two leaves in the FOV, one (above) treated with nPAA-MnO_2_ (drop applied) and one (below) untreated (SI Video S5). G). Two leaves in the FOV, one (above) infiltrated with nPAA-MnO_2_ and one (below) untreated (SI Video S6). Scan details are described in SI Table S3.

In additional experiments, drop application was tested to evaluate whether uptake could be followed inside the leaf. While Au NPs could be clearly detected on the surface of the leaf (Fig. 4C), no signs of uptake through stomata or other pathways were observed. Notably, in this experiment, the Au NP suspension was not amended with Silwet Gold, meaning it had a high surface tension, and this has been shown to significantly reduce NP uptake in barley leaves (Frank et al. 2025). Notably also, even if the NPs entered the leaf, they would be difficult to detect if they did not aggregate.

Iohexol, previously used to track foliar water films with synchrotron-based microCT (Frank et al. 2025), was tested here to evaluate the capabilities of lab-based microCT. In this experiment, the iohexol solution was applied as droplets onto a horizontally positioned barley leaf to monitor natural uptake, with a control leaf maintained in the same FOV. A scan was taken before droplet application and then continuously for 22 hours after droplet application. The time-series clearly showed that the iohexol remained detectable on the treated surface during the entire experiment, but some of it also penetrated quickly into the tissue. This is visible in a volume rendering of a scan taken after 18 h (Figure 4D). The time dependent distribution of iohexol can also be tracked by comparison of the same CT slice across different time scans as shown in Figure 4E, with close-ups highlighting volumetric changes in iohexol distribution with time. Zones of penetration of the contrast agent indicate uptake through open stomata. Overall, these results demonstrate that lab-based microCT can track the spatiotemporal dynamics of foliar uptake, allowing direct visualization of penetration sites and subsequent redistribution within the tissue.

In the next sets of experiments, drop application and infiltration tests were also conducted with potential nanoparticulate fertilizers (i.e., nHAp and nPAA-MnO_2_) intended for foliar application (Pinna et al. 2025, Minutello et al. 2025). However, the significantly lower contrast of these materials, and the fact they did not sufficiently aggregate, made visualization of these NPs within the tissue more challenging, particularly for HAp NPs. PAA-MnO_2_ NPs showed the highest potential, being clearly visible on the leaf surface (Figure 4F), but detection within the leaf tissue was challenging due to water-related artifacts in the pore network that could mimic NP accumulation (Figure 4G).

The nanofertilizers tested here were not forced to aggregate and further development is needed to visualize them, especially when using the drop application method. Overall, in our experiments, both drop application and direct infiltration allowed visualization of high contrast agents inside the tissue, appearing as localized high-intensity regions compared to control leaf tissues. In future studies, analyzing variations in high-intensity zones should be done systematically to provide quantitative assessments of foliar uptake from lab datasets, thereby opening perspectives for comparing delivery strategies and linking compound distribution to physiological responses.

## Conclusions

The methodology developed in this study enables micrometer-scale 4D X-ray lab microscopy of living leaf tissue, addressing the significant challenge of observing delicate plant structures under natural, undisturbed conditions. Maintaining plant physiological viability during prolonged X-ray exposures was key, achieved effectively using the horizontal sample setup, which allowed for time-series measurements extending beyond 20 hours. This optimized lab-based microCT system demonstrated the capacity to resolve cellular structures with a 1 µm pixel size. The method proved capable of tracking the uptake and redistribution of foliar-applied compounds, including contrast agents like iohexol, within the living tissue. These advancements open the window for widespread, accessible in situ 4D monitoring of physiological and anatomical processes in plants using lab-based microCT. Key challenges remain for studies employing nanomaterials with low-contrast, such as organic nanoparticles. Furthermore, since individual NPs are below the microCT resolution, detection necessitates their aggregation into micrometer-scale clusters. Future development must therefore prioritize overcoming resolution limits with multiscale investigations to track potential nanofertilizers (like HAp and PAA-MnO_2_ NPs). Equally important is assessing the tolerance of different plant species to extended scanning durations beyond 22 hours, to determine how long they can withstand radiation exposure without physiological impairment and to elucidate interspecific variability in their responses. These developments are crucial to fully leverage lab-based microCT for robustly comparing nutrient delivery systems and linking compound distribution to physiological responses.

## Acknowledgements

We thank the Danish Agency for Science Technology and Innovation for funding the instrument center DanScatt and the Danish National Facility for Imaging with X-rays, DANFIX. We thank Søren Nimb from the fine-mechanic workshop at DTU for support with the sample holder construction. This research was funded by the Novo Nordisk Foundation (grant no. NNF21OC0066114).

## Author contributions

F. Siracusa: data curation, investigation, formal analysis, methodology, software, writing – original draft

E.V. Kristensen: data curation, formal analysis, software, writing – review & editing

A. Szameitat: investigation, data curation, formal analysis, writing – review & editing

D. J. Tobler: conceptualization, supervision, validation, visualization, writing – original draft

I. Ertem: investigation, data curation, formal analysis

M. Frank: investigation, formal analysis, writing – review & editing

C. Wu: investigation, data curation, formal analysis, software, writing – review & editing

R. Abdurrahmanoglu: investigation, data curation, formal analysis, writing – review & editing

A. Pinna: investigation, data curation, formal analysis, writing – review & editing

F. Minutello: investigation, data curation, formal analysis, writing – review & editing

S. Husted: funding acquisition, project administration, resources, supervision, writing – review & editing

J.-C. Grivel:, resources, validation

R. Mokso: conceptualization, supervision, resources, funding acquisition, writing – review & editing

